# MR-μScope: a multi-scale and simultaneous MRI-miniaturized microscopy system for elucidating neurovascular coupling

**DOI:** 10.64898/2026.01.05.697821

**Authors:** Xinyi Zhu, Rui Li, Hui Li, Liangtao Gu, Liang Chen, Jiacheng Xu, Jingyin Chen, Ye Li, Markus Rudin, Garth Thompson, Ning Zhou, Wuwei Ren

## Abstract

Blood Oxygen Level Dependent functional MRI (BOLD-fMRI) revolutionized non-invasive brain mapping, yet the physiological origins of its signal, linking microscopic neurovascular activity to macroscopic hemodynamics, remain elusive. To bridge this gap, we developed MR-μScope: a simultaneous and multimodal platform that integrates preclinic high-field MRI (9.4T) and miniaturized fluorescence microscopy (Miniscope). Our system contains an MR-compatible Miniscope with electromagnetic shielding, a customized radiofrequency coil with a central optical window, and an adjustable animal cradle, enabling artifact-free acquisition of whole-brain fMRI alongside microscopic vascular dynamics. Phantom validation confirmed negligible cross-modal interference (fMRI SNR ≥ 15 dB; optical SNR ≥ 26 dB). MR-μScope captured stimulus-evoked microvascular dilation and blood flow velocity changes in the somatosensory cortex concurrent with BOLD signals, revealing vessel-size-dependent neurovascular coupling. MR-μScope provides an effective solution for deciphering multi-scale neurovascular interactions and offers unique potential for advancing research into brain function and disease mechanisms involving vascular pathology.

## 1. Introduction

Functional magnetic resonance imaging (fMRI) is a popular neuroimaging technique that maps brain activities in humans and animals. Owing to its unique advantages of non-invasiveness, balanced spatiotemporal resolution, and whole-brain coverage[1], fMRI has found a wide range of applications in characterizing functional architecture of the brain in healthy subjects and under pathological conditions such as stroke[2], neurodegenerative diseases[3], and psychiatry[4]. More recently fMRI has been used in the context of brain-computer interfaces[5]. Despite over three decades of extensive use, questions regarding the physiological origin of the Blood Oxygen Level Dependent (BOLD) fMRI signal, reflecting alterations in local cerebral blood flow and oxygenation, remain. It is not fully understood how neuronal activity within cellular networks is propagated to macroscopic BOLD signals, a complex process termed neurovascular coupling (NVC)[6]. Moreover, there is compelling evidence that cells different from neurons contribute to the observed fMRI signal changes including astrocytes[7], endothelial cells[8], and capillary pericytes[9]. These limitations are concerning because brain diseases are often accompanied by vascular dysfunction, including reduced blood vessel responsiveness, disrupted blood flow, and/or small vessel damage. As a result, altered fMRI responses might indicated alterations in neuronal function and/or the efficiency of NVC[10–12]. Given the multifaceted contributors to the BOLD effect, there is a critical need to record microscopic neural activity and hemodynamic changes alongside the macroscopic BOLD signals.

Compared with MRI, optical recording offers superior sensitivity and spatiotemporal resolution, and are capable of capturing subtle neural activities and hemodynamic changes, thus complementing fMRI effectively. For example, calcium imaging based on genetically encoded Ca²⁺ indicators enables mesoscale observation of activated neural populations[13] and microscale monitoring of individual cell dynamics[14]. Optical intrinsic signal imaging (OISI) exploits the intrinsic spectral properties of blood cells and tracks the fluctuations of blood flow and oxygenation at multiple scales[15]. Laser Doppler flowmetry and laser speckle contrast imaging are mostly used to quantify cerebral hemodynamics across the whole brain in rodents[16]. With the richness of optical imaging and sensing techniques, synergizing the image contrast of optics and MRI at varied spatial scales would allow researchers to understand the full process of how microscopic neural activity drives quantitative microscopic hemodynamics, which subsequently integrates into the non-quantitative macroscopic BOLD signal.

A sequential dual-modal strategy combining two imaging modalities is highly practical and has been effectively adopted to study the hemodynamic changes and BOLD signals[6, 17, 18]. Nevertheless, capturing the spatiotemporal signal dynamics within one subject requires a hybrid system enabling simultaneous recording of MRI and optical signals. The combination of fiber-photometry and MRI has been widely adopted in mechanistic studies of BOLD contrast[19–21]. While offering high speed and sensitivity, fiber-photometry lacks spatial resolution. By replacing a single fiber with a fiber bundle spatially resolved optical signals could be recovered, enabling simultaneous calcium and fMRI signal recording across the entire cortex[22, 23]. The resulting spatial resolution of calcium imaging acquired by a fiber bundle is determined by the dimension of individual fibers, which is up to tens of micrometers, far from resolving individual brain cells or microvasculature. Other concurrent MRI-optics systems combining photoacoustic imaging[24], OISI[25], diffuse optical imaging[26], and Laser Doppler[27] have also been proposed. To date, most reported concurrent MRI-optics systems focus on the macroscopic or mesoscopic optical readout. Development costs for such systems are typically high as they are based on expensive optical components such as fiber bundles[22, 23]. Yet, for investigating the link between the neural activities/vascular responses at the microscopic level and the hemodynamic adaptations/BOLD signals at the macroscopic level, a more affordable and robust MRI-optics hybrid system covering multiple scales would be highly attractive[28].

Here we present the MR-μScope, a simultaneous and multiscale dual-imaging system that combines miniaturized fluorescence microscopy (Miniscope) and high-field MRI, featuring high optical spatial resolution (∼0.7 μm), fast optical readout (60 Hz), whole-brain fMRI coverage for mice (FOV ≥ 15 × 15 × 10 mm³) and low cost. To ensure dual imaging quality, we developed an MR-compatible Miniscope, a custom transceiver-integrated coil, and a custom animal cradle. After phantom validation, MR-μScope acquired synchronized measurement of cortical microvascular activities and whole-brain fMRI in mice for the first time. Multiparametric information including local vascular diameter, blood flow velocity, and BOLD signals in the cortex was extracted from the raw data, along with correlation analysis. Furthermore, our system recorded for the first time the differential contributions of various-sized vessels (15-60 μm) to NVC. These findings revealed activity induced vascular changes in individual vessels, enabling a direct comparison between microvascular events and global hemodynamic responses. In short, MR-μScope bridges the gap between microscopic and macroscopic imaging, offering new opportunities for multimodal, multiscale brain research in awake and anesthetized animals.

## 2. Results

### 2.1 Hybrid system for BOLD fMRI and microscopic imaging

We have devised the MR-μScope, a hybrid imaging system incorporating an MRI-compatible miniaturized fluorescence microscope, a customized radiofrequency (RF) coil, an animal cradle, and physiological-monitoring/maintenance equipment, fully integrated to a preclinical 9.4T-MRI scanner (BioSpec 94/30, Bruker Biospin GmbH, Germany) (Fig. 1a). The Miniscope part is designed as an MR-compatible insert, which is adapted from the open-source UCLA Miniscope [29] and allows for noise-free microscopic imaging in the presence of high magnetic field up to 9.4T. The customized RF coil is compatible with the optical hardware and enables synchronized acquisition of whole-brain structural/functional MRI with a field-of-view (FOV) of 15 × 15 × 10 mm³. For the optical readout, we achieved a spatial resolution of ∼0.7 μm and a frame rate of 60 Hz.

**Fig. 1.**
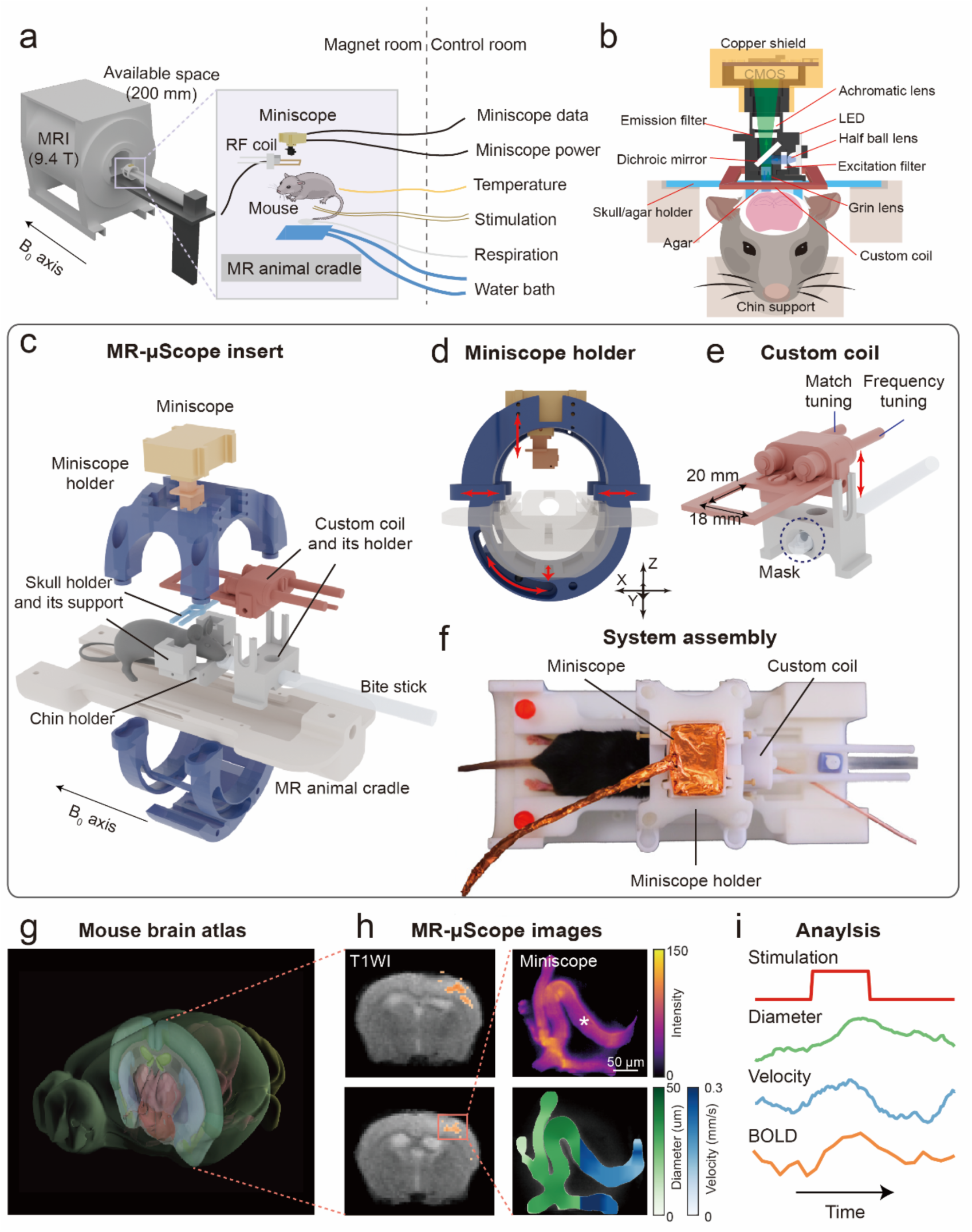
The experimental set-up of MR-μScope, an MRI-miniaturized microscopy hybrid system. (a) A schematic of MR-μScope based on a 9.4 T MR scanner, into which the MR-compatible Miniscope, the custom-made RF coil, the animal cradle, the animal, and the physiological-monitoring/maintenance equipment. (b) Detailed design and installation of the Miniscope, which positioned above the mouse brain with the GRIN lens passing through the window of the custom-made RF surface coil allowing adjustment to optimal adjustment of working distance through the custom surface coil. (c) Exploded-view drawing of the MR-μScope main body, consisting of the Miniscope, the RF coil, the skull holder and the cradle. (d) The adaptive Miniscope holder allows 4 degrees-of-freedom (DOFs) of movement assuring robust imaging performance with optimal positioning and orientation of the heads of individual mice. (e) Detailed design of the custom-made RF coil and its holder. The coil holder contains the anesthesia mask, and the bite stick for head fixation. (f) Top view photo of the MR-μScope insert. (g) A 3D reference atlas of the mouse brain generated from Allen Mouse Brain Common Coordinate Framework[52]. (h) Two adjacent coronal BOLD-fMRI images during unilateral hindpaw stimulation overlaid on T1-weighted (T1W) images cross-sectional anatomical reference images (left) and fluorescence microscopic images of the cerebral microvasculature used to derive morphological (vessel diameter) and functional (blood flow velocity) information (right). (i) Quantitative analysis: Stimulus-evoked (red trace) changes in average BOLD signal amplitude (orange), as well as vessel diameter (green) and blood flow velocity (blue) for a selected microvessel (*) labeled in (h)

To prevent interference between the two imaging modalities, we optimized the Miniscope design in several aspects (Fig. 1b). Firstly, the CMOS imaging sensor PCB was re-designed by replacing the ferromagnetic components to warrant MR compatibility. The active metal crystal oscillator in the clock module was replaced with a passive ceramic crystal oscillator, while an additional low-dropout regulator (LDO) was used to replace the ferrite beads of the original DC-DC module for maintaining sufficient power supply (Supple. Fig. 1). Secondly, a custom enclosure made from copper was used for electromagnetic shielding to minimize the interference from B0/B1 fields, and a LED PCB was further employed to reduce signal losses. Lastly, we developed a radiofrequency (RF) customized transceiver surface coil with a 20 × 18 mm² window (Fig. 1e) to maintain fMRI signal-to-noise ratio (SNR) at ∼15 dB during simultaneous multimodal imaging, showing no significant difference in SNR when comparing the hybrid setup with the MRI alone (Fig. 2e).

**Fig. 2.**
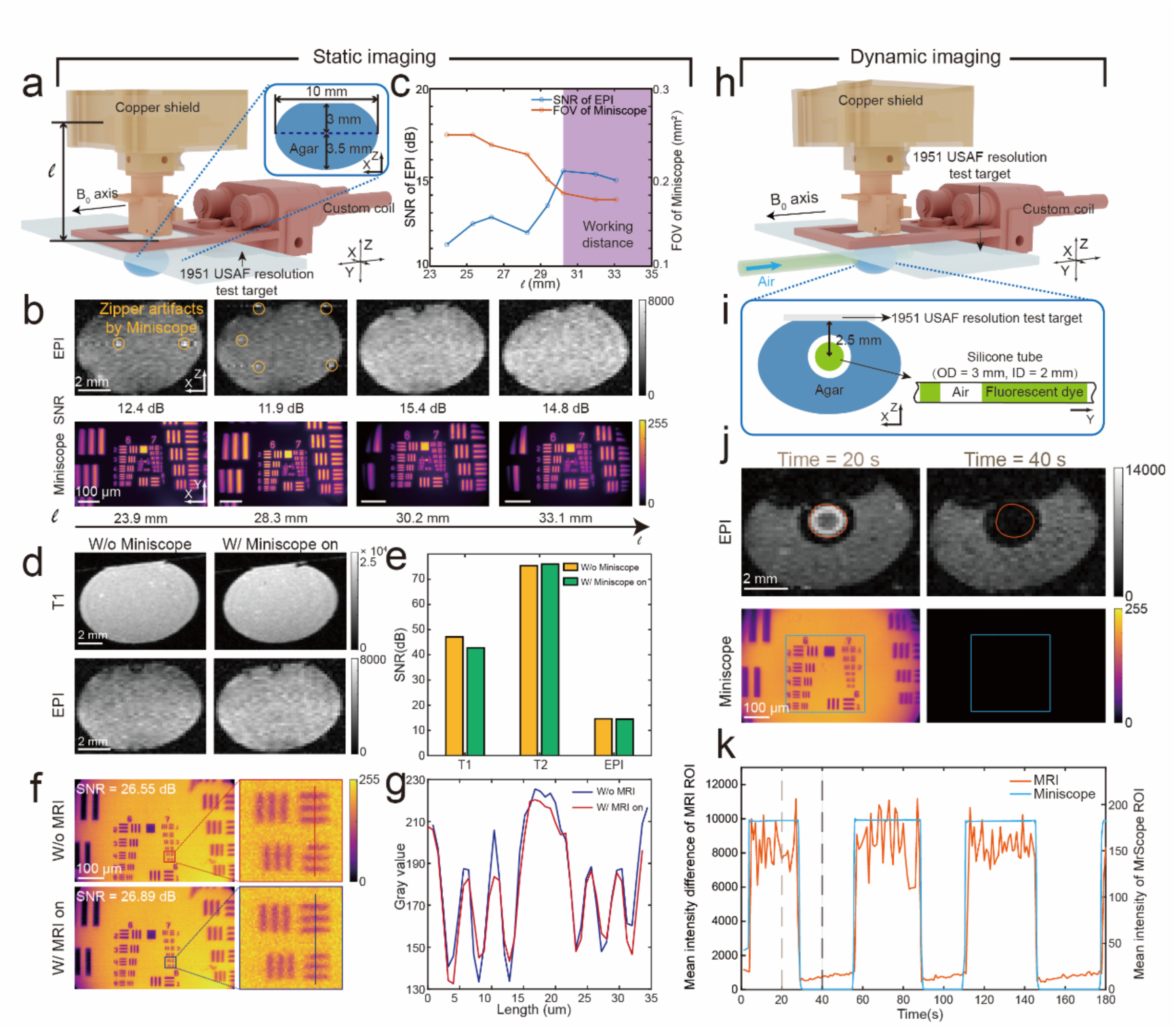
Evaluation of system performance with regard to sensitivity and interference using phantom samples under static (a-g) and dynamic (h-k) conditions. (a) Test setup for static imaging with a 1951 USAF resolution board and an agar-made ellipsoid placed underneath the RF coil. Images were acquired for different working distances l as indicated in the figure. (b) EPI-MRI and Miniscope images as a function of l. EPI images show reduced zipper artifacts (yellow circles) and improved SNR-values, while the FOV of Miniscope images becomes smaller. (c) Quantitative analysis: In the range of 30<l<33 mm, a close to maximum SNR for EPI-MRI was achieved at the expense of a slightly reduced FOV of ∼ 4 × 4 mm2, considered as the optimal working distance (the purpleregion). (d) Qualitative appearance of T1W and EPI-MRI images with Miniscope switched on (left) and off (right) for a working distance l of 30.2 mm. (e) Quantitative analysis of SNR of T1-, T2-weighted and EPI images with and without Miniscope shows no or minimal degradation of MRI signals with Miniscope on. (f) Miniscope images obtained with and without MRI in operation. (g) Line intensity profiles in Miniscope images in (f) reveal minimal degradation of the optical signal by MRI operation. (h, i) The phantom experiment for dynamic imaging with the same resolution board and an agar-made ellipsoid containing a silicone tube (outer/inner diameter: 3/2 mm) filled with solution with fluorescence dye in stereo (h) and cross-sectional view (i). During the measurement, the solution containing the dye flows through the agar in y direction. The solution column in the tube is interrupted with air bubbles. (j) EPI-MRI and Miniscope images at t = 20 s and 40 s (see also Fig.(k)). When the solution containing the fluorescence dye flows right underneath the Miniscope lenses (t = 20 s), the illuminated resolution board is detected by Miniscope and a bright ring structure is observed in the MRI image. When an air bubble appears in the FOV, neither signal is observed. (k) Time lapse of the mean intensity values of ROIs in both EPI-MRI and Miniscope images as indicated in (j).

To achieve robust in vivo imaging performance, a custom mouse cradle equipped with adjustable holders was designed to provide flexible positioning and immobilization inside a 114 mm-diameter MRI bore. A dual-pin skull holder secures the mouse head to the cradle, together with the jaw holder and a breathing mask (Fig. 1b, c). A ring-shaped Miniscope holder enables movement of the optical device along 4 degrees of freedom (DOF) to compensate for inter-animal variations in cranial window positioning (Fig. 1d). The height-adjustable coil is positioned above the mouse head with the Miniscope gradient-index (GRIN) lens passing through its window (Fig. 1e). Furthermore, the animal cradle integrates temperature control, respiratory gating, and electrical stimulation interfaces to support in-vivo task-based experiments, permitting simultaneous acquisition of whole-brain fMRI and localized vascular fluorescence signals in response to stimulation inside the MRI scanner (Fig. 1a). A photograph of system assembly is shown in Fig. 1f. Detailed system description refers to online Methods and Supple. Fig. 4.

To demonstrate the performance of the MR-compatible Miniscope with a custom-designed MRI surface coil under *in vivo* conditions, we acquired structural MRI images of the mouse brain, overlaid with stimulation-evoked BOLD-fMRI maps, and simultaneously recorded fluorescence microscopy images of localized micro-vessels in targeted brain areas (Fig. 1h). These multimodal time-resolved images yield both morphological and hemodynamic information such as the transient changes of the vascular diameter, blood flow velocity, and BOLD contrast in response stimulation. Analysis of the synchronized BOLD-fMRI and fluorescence signals enables the investigation of the relationship between local vascular adaptations and whole-brain hemodynamics (Fig. 1i).

### 2.2 Characterization and compatibility test based on phantom experiments

#### 1) Static imaging experiment

Before in vivo studies, the system performance was extensively evaluated using two phantoms. Initially, we designed a static imaging experiment to evaluate the image quality acquired in dual-mode operation with metrics such as SNR and spatial resolution (Fig. 2, the left panel). A brain-mimicking ellipsoid phantom (Phantom A, see online Methods) made of agar (dimension, lengths on x-/y-/z-axis = 10/10/7 mm) was housed in a 3D-printed casing and placed underneath the RF coil for MRI recording. The lens assembly of Miniscope was mounted through the coil opening such that the GRIN lens (#64526, Edmund Optics, USA; diameter, 1.8 mm; length, 4.31 mm; focal length, 0.23 mm) was pressed against the positive contrast 1951 USAF resolution board (R3L1S4P1, Thorlabs, New Jersey, USA) (Fig. 2a).

We firstly optimized the working distance ℓ between the sensor plane and the coil surface to ensure high image quality of both MRI and optical images. We selected 8 positions ranging from ℓ = 23.9 mm to ℓ = 33.1 mm, covering the entire adjustable focus range of the Miniscope, to evaluate the effect of the working Miniscope on MR images using an echo-planar imaging (EPI) sequence. For ℓ < 30.2 mm, the EPI images exhibit strong zipper artifacts, which are not observed anymore for ℓ ≥ 30.2 mm. SNR values of EPI images are of the order of 11 to 12 dB for ℓ ≥ 28.3 mm, and increase steadily for a large distance reaching a plateau of 15 dB for ℓ ≥ 30.2 mm (Fig. 2c). Simultaneously, we recorded optical images by tuning the position of the achromatic lens and assessed the FOV and spatial resolution for each image at different ℓ values (Fig. 2c). As ℓ increases from 23.9 mm to 33.1 mm, Miniscope FOV decreases from 0.62 × 0.40 mm² to 0.51 × 0.33 mm², while the spatial resolution improves from 0.83 μm to 0.68 μm (Fig. 2c). These data indicate an optimal working distance in the range of ℓ = 30.2 mm (FOV: 0.53 × 0.34 mm²; resolution: 0.71 μm) to ℓ = 33.1 mm (FOV: 0.51 × 0.33 mm²; resolution: 0.68 μm).

Next, we evaluated the interference of an operating MR-μScope on MRI performance for various MRI acquisition sequences: T1-weighted images (T1WIs) using a fast low-angle shot (FLASH) sequence, T2-weighted images (T2WIs) using a rapid acquisition with relaxation enhancement (RARE) sequence, and EPI sequence. The results show that the simultaneous operation of the Miniscope at an optimal working distance inside the MRI scanner did not significantly decrease the quality of MR images as reflected by the SNR values: SNR of T1WI/T2WI/EPI ≥ 40/70/12 dB (Fig. 2d, e). The simultaneously recorded optical images also exhibit a high SNR value of up to 26.89 dB, identical within error tolerance when compared with the measurement taken outside the MRI scanner (26.55 dB) (Fig. 2f, g).

#### 2) Dynamic imaging experiment

We took a further step to evaluate the performance of MR-μScope under dynamic conditions (flow). Similar to the previous experiment, an ellipsoid agar phantom (Phantom B, see online Methods) and a 1951 USAF resolution board were placed below the coil and GRIN lens (Fig. 2h) at the optimal working distance as previously defined. A silicone tube (inner diameter, 2 mm) was embedded within the agar sample, through which a fluorescein isothiocyanate-dextran aqueous solution (1 mg/mL) was circulated at a velocity of ∼3.3 mm/s (Fig. 2i). The solution column contained air bubbles allowing the modulation of the signals associated to the flowing medium. EPI-MRI was performed, with the Miniscope simultaneously operating in the scanner. We observed that the appearance of the fluorescence dye within the FOV renders the microscopic resolution board visible to Miniscope. Simultaneously, EPI-MRI captures the flowing water solution column in the silicone tube, which appears as a bright structure with a dark ring (silicone wall of the tube) surrounded by agar signal (Fig. 2j). In contrast, both Miniscope and tube-associated MRI signals disappear if an air bubble passes underneath the RF coil and GRIN lens (Fig. 2k). The relatively high flow velocity of the fluorescent dye (∼3.3 mm/s), with the fastest flow occurring in the vascular central region (assuming a laminar flow profile), leads to the characterstic ring enhancement observed. Unsaturated spins enter the imaging plane in between subsequent excitation, yielding the signal enhancement observed. Flow at the center of the tube is so high that these unsaturated water proton spins leave the imaging plane in between excitation and detection, which explains the reduced signal intensity at the center of the tube as compared to regions closer to the tube wall, where the flow velocity is lower and no outflow occurs (Fig. 2j). We have demonstrated that our system is capable of acquiring high-fidelity and synchronized MRI/optical images, paving the road for in vivo biological studies.

### 2.3 Evoked BOLD and microvascular responses

Following the characterization of the system in phantom studies, we performed in vivo mouse brain imaging, by assessing microscopic vascular dynamics and BOLD signal variations simultaneously in the somatosensory cortex during unilateral hind-paw stimulation. Prior to data acquisition, we performed a cranial window surgery over the right cortical S1HL region (hindlimb region of the primary somatosensory cortex) of the mouse, which receives somatosensory input from the hindlimb and can be used to observe the relationship between cortical BOLD signals and changes in vascular diameter and blood flow velocity, and implanted a GRIN lens (mentioned in 2.2). The lens was securely fixed using a 3D-printed, resin-cast dual-pin skull holder and dental cement, with the remaining cavity filled with agarose gel to eliminate air gaps that could potentially cause MRI artifacts. For enhancement of the microvasculature, fluorescence dye FITC-Dextran (0.3 mL, 3 mg/mL) was injected intravenously into the tail vein. The details of animal imaging studies are given in online Methods.

Following the simultaneous acquisition of BOLD-fMRI and fluorescence microscopic images during the stimulation paradigm, images were analyzed as follows (the workflow of multimodal data processing is depicted in Supple. Fig. 5). Fluorescence image data were further processed to extract information about mean microvascular diameter and blood flow velocity (Fig. 3a). Both the mean diameter/velocity maps and temporal response variations from individual mice demonstrate that the cortical pial vessels in S1HL exhibited dilation and increased flow velocity in response to electrical stimulation, with response timing synchronized to the stimulation cycle (Fig. 3b). Simultaneous fMRI recordings revealed transient BOLD changes with the primary somatosensory cortex (S1) (Fig. 3c). The individual temporal fluctuations of the BOLD intensity show that the electrical stimulation can specifically trigger signal enhancement in the target brain area (Fig. 3d). The delay of response onset time relative to the stimulus onset are 1.64 ± 1.67 s and 0.67 ± 1.20 s for diameter dilation and flow velocity, respectively, and 4.47 ± 2.94 s slightly longer for the BOLD transient. The time to peak of diameter and velocity changes are 8.29 ± 2.00 s and 7.98 ± 1.68 s, both shorter than that of BOLD transient, 10.58 ± 2.39 s.

**Fig. 3.**
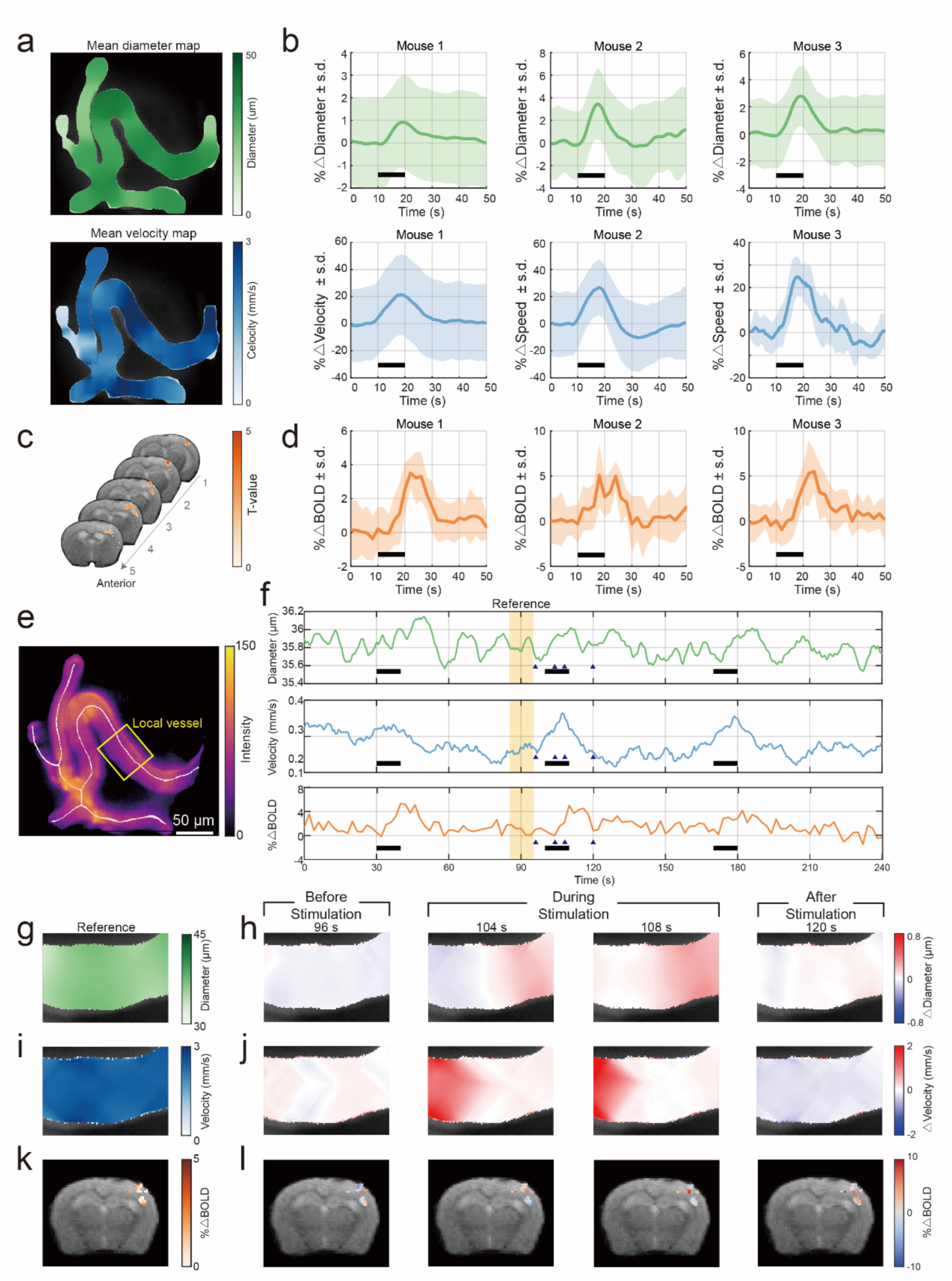
Microscopic recordings of vascular dynamics and macroscopic BOLD signal variations evoked by unilateral hind-paw stimulation. (a) The mean-value maps of microvascular diameter (upper) and velocity (lower) for a reference mouse during rest. (b) Quantitative analysis of changes in microvascular diameter (green) and velocity (blue) changes for 3 example mice averaged over 5 subsequent block stimulation periods. The duration of a stimulation block was 10 s indicated by the black line. The shaded area represents the standard deviation (SD). (c) fMRI data for the mouse shown in (a) with activated areas obtained from standard generalized linear modeling (p<0.05). (d) Averaged BOLD changes for the mice shown in (b). (e) Extracted skeleton of vessels in the mouse shown in (a). Results of local vascular changes in a yellow box are shown in (h-l). (f) Evoked microvascular responses in diameter and velocity are compared with the whole-brain BOLD responses for 3 subsequent stimulation blocks. For the quantitative analysis, data in the resting interval t = 85-95 s were averaged and used as the reference images (IRef) in (g), (i), (k). Difference images I(t)-IRef of microvascular diameter (h), velocity (j) and BOLD (l) signal at 4 time points are compared (t =86, 104, 108 and 120 s, marked with triangles in (f)). All these time points correspond to the moments before, during and after the stimulus.

To better understand these synergetic responses at both microscopic and macroscopic levels, we compared the transient responses of microvascular dilation and blood flow changes in a local vessel of the right S1HL region, and the BOLD signal changes in the corresponding S1 region during 3 consecutive stimuli in an identical mouse (Fig. 3e). With reference to a pre-stimulation baseline image (averaged over the interval 85 ≤ t ≤ 95 s, Fig. 3g, i, k), we show the difference images of local microvascular response mapping and macroscopic BOLD changes at three stages of a single trial, i.e., immediately before (96 s), during (104, 108 s), and after stimulation (120 s; Fig. 3h, j, l). Strong positive signals are observed during stimulation (t = 104, 108 s), indicating substantial activation at S1. After stimulation cessation (t = 120 s), the overall intensity tends to return to a baseline (Fig. 3h, j, l). Supple. Video shows the recorded Miniscope and EPI images during one stimulus, and the local temporal variations of the three parameters. The results also suggest that the changes in microvascular diameter and flow velocity precede alterations in the BOLD signal (Fig. 3b, d), indicating that local hemodynamic responses form the physiological basis for subsequent BOLD activation in functional MRI, consistent with the previous study using two-photon microscopy and fMRI separately[30].

### 2.4 Dependence of evoked responses on stimulation frequency

Multi-scale readouts of vascular responses to peripheral electrical stimuli were evaluated for three different stimulation frequencies of 3 Hz, 6 Hz, and 9 Hz, respectively, while the remaining parameters of the stimulation are constant: current strength at 1 mA, duty cycle at 9%, duration of stimulation block at 10 s with a resting interval of 60 s. Five mice were studied. Evoked microvascular responses and whole-brain BOLD-fMRI signals are shown for an example mouse at varying stimulation frequencies (Fig. 4). A raw fluorescence image of cortical pial vessels is displayed with four segments labeled (Fig. 4a). Stimulus evoked BOLD-fMRI signals were observed in the cortical S1 for all three stimulation frequencies (Fig. 4b). For clear visualization and analysis purposes, stimulus-induced changes in the microvascular parameters ‘vessel diameter’ (Fig. 4c) and ‘flow velocity’ (Fig. 4d) of the four vascular segments labelled are plotted alongside with the BOLD signal variations (Fig. 4e), with the stimulation signal sequence shown at the top of Fig. 4c. The selected vascular segments exhibit stronger diameter dilation at high-frequency stimulation with 1.6 ±1.0 % at 3 Hz (mean ± S.D.), 2.8 ± 0.5% at 6 Hz, and 3.1 ± 1.0% at 9 Hz (Fig. 4c). In comparison, flow velocity traces fluctuate at all three frequencies with percentage changes of 25.8 ± 10.3% at 3 Hz, 25.0 ± 10.9% at 6 Hz, and 27.7 ± 9.4% at 9 Hz, showing a similar increasing pattern as the diameter change as the frequency rises (Fig. 4d). Similarly, increased evoked BOLD signals are observed for high frequencies with the percentage changes 1.3 ± 0.6% at 3 Hz, 1.9 ± 0.8 % at 6 Hz, and 2.0 ± 0.9 % at 9 Hz, respectively.

**Fig. 4.**
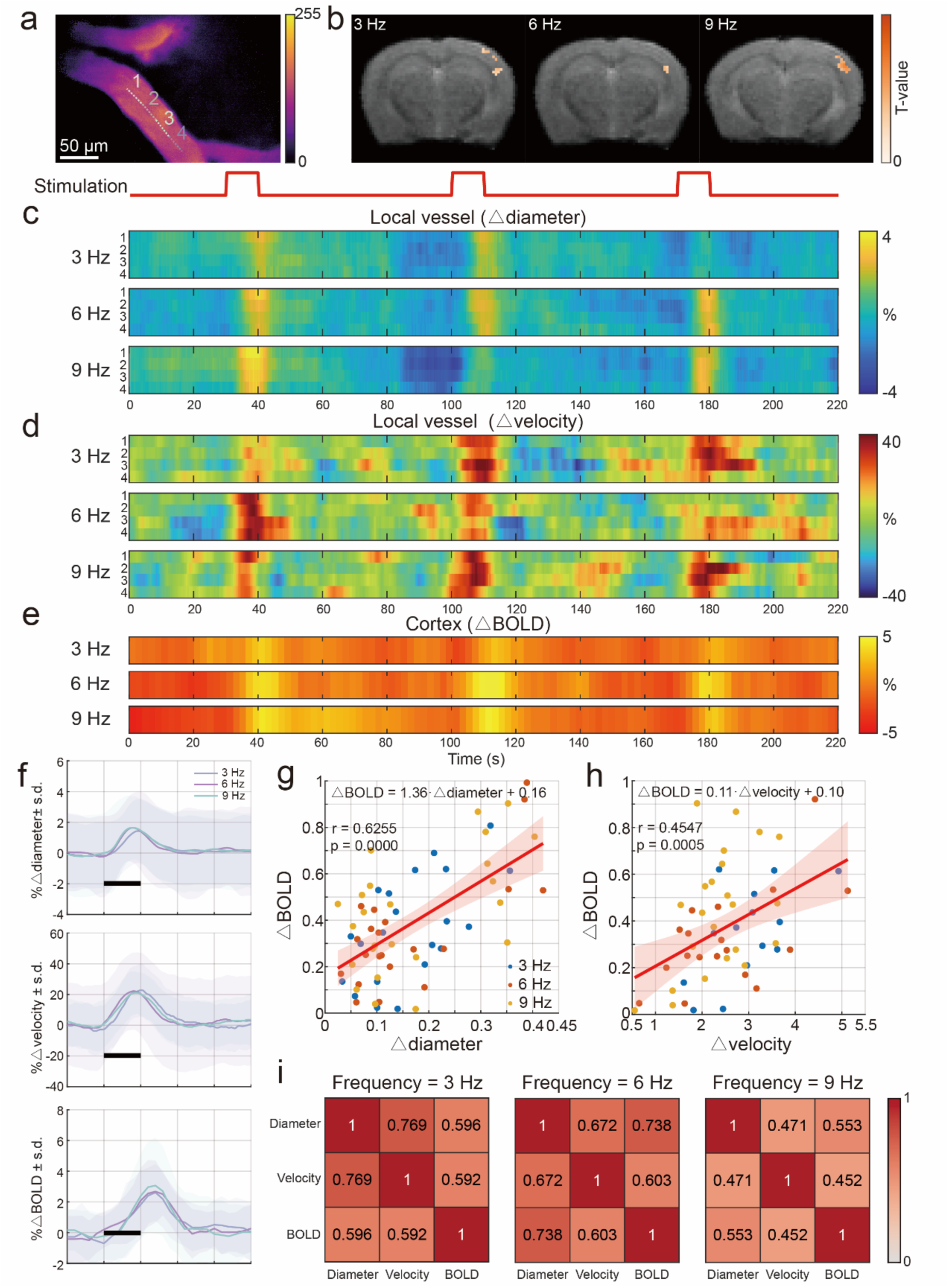
Evoked microvascular responses and whole-brain BOLD-fMRI signals as a function of stimulation frequencies. (a) Microscopic vessels in an example mouse brain acquired by Miniscope. (b) fMRI ROI data for the same mouse stimulation frequencies 3 Hz, 6 Hz and 9 Hz. ROIs for quantitative analysis were determined using the standard generalized linear model. (c) Stimulus-evoked diameter changes of vascular segments 1-4 (labeled in a) for 3 subsequent stimulation blocks of 10 s with frequencies 3, 6, and 9 Hz (red trace). Note weaker response amplitude for 3Hz as compared to 6Hz and 9Hz stimulation and decline of response amplitude for subsequent stimulations, indicating adaptation. (d) Stimulus-evoked velocity changes of these microvascular segments. Note significantly higher fluctuations during the resting periods which impair the quantitative analysis. Again, stimulation frequencies 6Hz and 9 Hz yield more reliable signals. (e) The simultaneously recorded fMRI changes in ROI yield robust signal responses showing the same qualitative behaviour as the diameter measures. (f) Averaged responses of diameter changes (upper), velocity changes (middle) and fMRI changes (lower) for the three stimulation frequencies across all 5 mice and all 5 stimulation blocks. The block duration is indicated by a black line. The SD across trials is represented by the shaded area. (g, h) Correlation of BOLD signal changes with diameter changes and velocity changes (calculated using area under the curve), resulted in r = 0.6255/p = 0.0000 and r = 0.4547/p = 0.0005, respectively. (i) Correlation heatmap of stimulus-evoked changes of diameter, velocity and BOLD at f = 3 Hz, 6 Hz and 9 Hz.

We further analyzed the data by averaging the vascular diameter/flow velocity changes, and BOLD signals across all trials from all five mice as a function of stimulation frequencies (Fig. 4f). In general, vascular diameter changes increase in response to stimulation, and become pronounced at 6 Hz and 9 Hz (1.4 ± 2.1% at 3 Hz, 1.6 ± 2.2% at 6 Hz, and 1.6 ± 2.1% at 9 Hz). Quantitative flow velocity across all mice regarding frequencies are similar but features a larger variation amplitude than diameter, with 22.8 ± 17.0% at 3 Hz, 22.2 ± 25.1% at 6 Hz, and 21.1 ± 13.6% at 9 Hz, whereas BOLD signals exhibit stronger responses at 9 Hz (3.1 ± 3.1% at 9 Hz) and are weaker at lower frequencies (2.6 ± 1.8% at 3 Hz and 2.7 ± 2.0% at 6 Hz).

For quantitative comparison, we parametrized the experimental time profiles of the three readouts (diameter, velocity, and BOLD signal) using gamma-variate function fitting. The overall response for each readout was estimated by area-under-the-curve (AUC) calculation. The correlation analysis between each hemodynamic parameter and the BOLD signal reveals a significant linear relationship (p < 0.05), with the diameter-BOLD correlation (Pearson correlation coefficient: r = 0.63) being stronger than the velocity-BOLD correlation (r = 0.45) (Fig. 4g-h). Frequency-specific analysis yields the lowest correlation at 9 Hz for both the diameter-BOLD pair (r = 0.55) and the velocity-BOLD pair (r = 0.45). At other frequencies, the diameter -BOLD correlation increases significantly (3 Hz: r = 0.60; 6 Hz: r = 0.74), while the velocity -BOLD remains essentially unchanged (3 Hz: r = 0.59; 6 Hz: r = 0.60) (Fig. 4i). Irrespective of the simulation frequency, the r-values of the diameter-BOLD correlation were found to be larger than those of the velocity-BOLD correlation, suggesting that vascular diameter changes may serve as a more reliable physiological predictor of BOLD signal variations in the neurovascular coupling process. A possible explanation is that the BOLD signals, which result from relative changes in deoxyhemoglobin, are largely determined by the total blood volume and, hence, the total amount of oxy-/deoxy-hemoglobin in the region of interest[31].

### 2.5 Contribution of microvascular sizes to BOLD-fMRI

To investigate the contribution of cortical microvascular size to the macroscopic BOLD-fMRI signals, we categorized all vascular segments into three groups according to the local mean diameter value throughout the stimulation experiments (small: 15-30 𝜇m, medium: 30-40 𝜇m, large: 40-60 𝜇m). Fig. 5a is a representative fluorescence image acquired by MR-μScope under 9 Hz stimulation, which contains multiple microscopic vessels of different sizes. Vascular segments from 3 categories are labeled with mean diameters of 22.9 𝜇m, 40.9 𝜇m, and 53.9 𝜇m, respectively. The corresponding macroscopic BOLD-fMRI images from the same trial are also shown (Fig. 5b). All vascular segments (vessel 1-3) exhibit diameter dilation during stimulation (Fig. 5c), most pronounced in small vessels but less obvious in medium- and large-sized micro-vessels (Δdiameter = 5.0 ± 16.3%, small; 0.9 ± 13.0% and 0.5 ± 4.3%, large). In comparison, flow velocity fluctuations show greater variability and consistently display stimulation-triggered responses (Fig. 5d), with the largest amplitude observed in small and medium-sized vessels (Δvelocity = 7.2 ± 7.6%, small; 7.5 ± 12.5%, medium; 4.0 ± 5.9%, large). The stimuli also induce significant BOLD signal increases (Fig.5e) in S1 cortex (ΔBOLD = 2.0 ± 0.9%) (Fig. 5c-e).

**Fig. 5.**
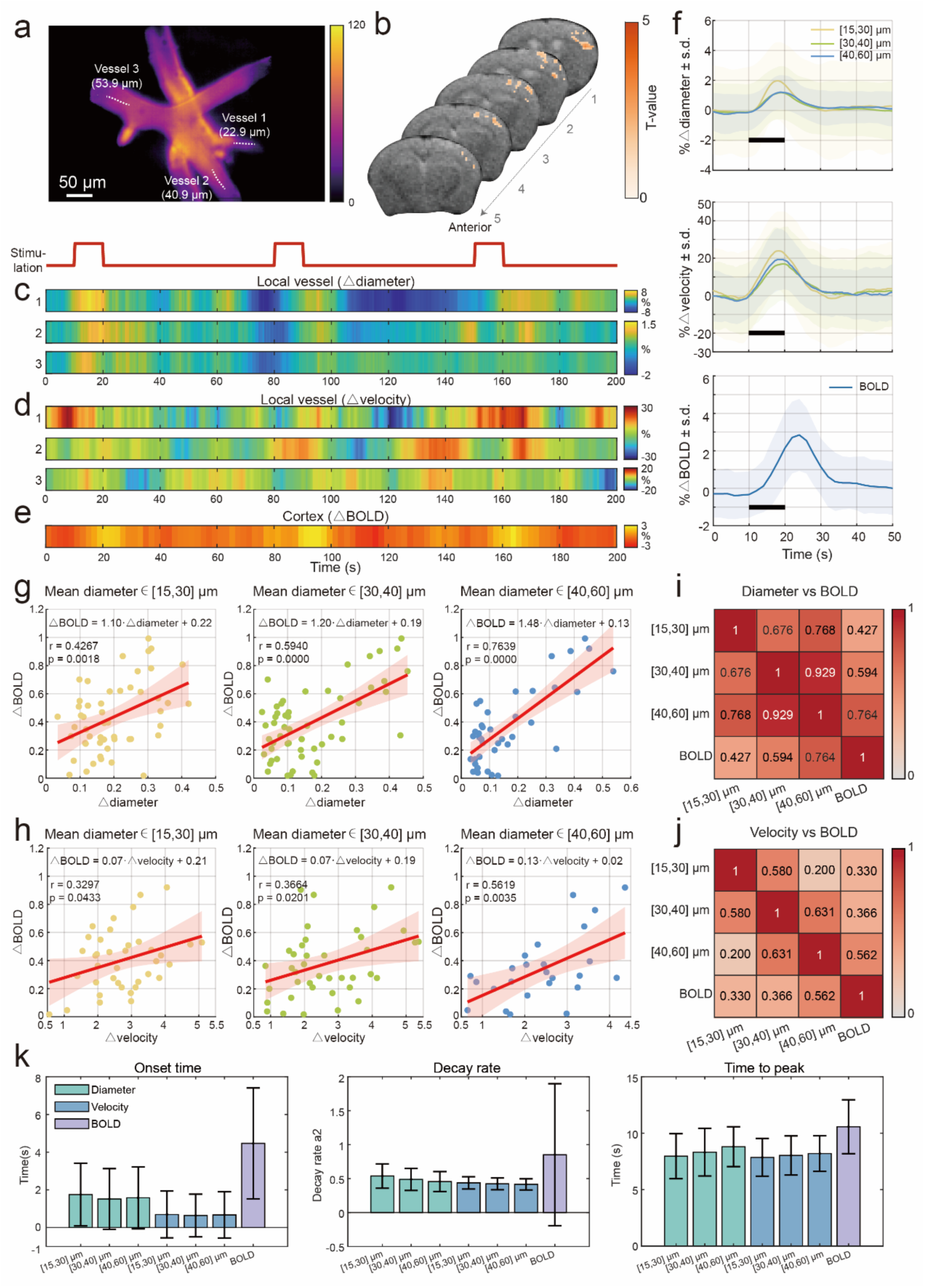
The dependence of vascular size for the evoked microvascular responses and fMRI signals. (a) Microvasculature in an example mouse imaged by Miniscope. 3 vessels of different sizes are marked with dashed lines and used for further analysis. (b) fMRI ROI data for the same example mouse. (c, d) Stimulus-evoked diameter (d) and velocity changes (c) in local vessels 1, 2 and 3 with stimulation frequency = 9 Hz responding to 3 stimulation blocks (red trace). (e) The simultaneously recorded changes in BOLD amplitude in ROI. (f) Analysis of responses of diameter changes (upper), velocity changes (middle) of vessels with different sizes and average fMRI changes (lower) averaged across all 5 mice and 5 stimulation blocks for 3 stimulation frequencies. The stimulation block is indicated by a black line. The SD across trials is represented by the shaded area. (g, h) Correlation of BOLD signal changes with diameter changes and velocity changes (calculated using area under the curve) for different vessel size groups, resulted in r = 0.4267, 0.5940, 0.7639/p = 0.0018, 0.0000, 0.0000, and r = 0.3297, 0.3664, 0.5619/p = 0.0433, 0.0201, 0.0035, respectively. (i) Correlation between microvascular diameter and BOLD classified by vessel size groups. (j) Correlation between microvascular velocity and BOLD classified by vessel size groups. (k) Comparison of the dynamic parameters (onset time, decay rate and time to peak) of diameter changes and velocity changes for different vessel sizes and BOLD signal changes.

We further averaged the vascular diameter/velocity changes within each size group (Fig. 5f). Similar to the trend of individual data, diameter change amplitudes show the greatest response. (Δdiameter: 2.0 ± 2.6%, small; 1.2 ± 1.8%, medium; 1.2 ± 1.1%, large), and so as the flow velocity change amplitudes (Δvelocity = 23.9 ± 20.9%, small; 17.0 ± 19.6%, medium; 19.3 ± 16.0%, large). Statistical analysis shows that BOLD signal variation changes 2.8 ± 1.9%.

Correlation analysis between vascular diameter/blood flow velocity and BOLD signals across different-sized blood vessels reveal a significant linear relationship, with increasing r-values as the vessel size increases (Fig. 5g, h) with r = 0.43 for small, r = 0.59 for medium sized, and r = 0.76 for large vessels for the diameter-BOLD and r = 0.33 for small, r = 0.37 for medium sized, and r = 0.56 for large vessels for the velocity BOLD correlation. This indicates differential contributions of microvascular hierarchies to the BOLD signal, where larger vessels appear to play a dominant. Significant inter-vessel correlations (Fig. 5i, j) are also observed for diameter changes (small-medium: r = 0.68; small-large: r = 0.77; medium-large: r = 0.93) and blood flow velocity across all size categories (small-medium: r = 0.58; small-large: r = 0.63; medium-large: r = 0.20). Previously reported modeling methods of the BOLD signal have indicated that the larger end of microvasculature drives the BOLD signal at the echo times used in our study[32]. Nevertheless, the specific diameter changes observed in this study differ from their simulation.

We further investigated the dynamic characteristics of the three parameters response onset time 𝑡_o_, decay rate 𝑘, and time-to-peak 𝑇𝑇𝑃 for diameter/velocity signals of different sized vessels and the BOsLD signals (Fig. 5k). For diameter changes, the small vessels feature the shortest onset time (small: 𝑡_o,d_ = 1.75 ± 1.66 s), with relatively small difference between medium and large vessels (medium: 1.51 ± 1.61 s; large: 1.58 ± 1.65 s). A similar trend was found in flow velocity changes (small: 𝑡_o,v_ = 0.70 ± 1.24 s; medium: 0.64 ± 1.13 s; large: 0.67 ± 1.23 s). The onset times of both vessel diameter and flow velocity are much shorter than that of the BOLD signal (𝑡_o,*BOLD*_= 4.47 ± 2.94 s). The decay rate of both diameter change (small: 𝑘_d_ = 0.54 ± 0.18; medium: 0.49 ± 0.16; large: 0.46 ± 0.15) and flow velocity change (small: 𝑘_v_ = 0.44 ± 0.09; medium: 0.42 ± 0.09; large: 0.41 ± 0.08) are similar appear to be rather insensitive to the vessel size. The decay rate of the BOLD signal was found to be larger (𝑘_*BOLD*_ = 0.85 ± 1.05). The time-to-peak of both diameter and flow velocity changes increase with increasing diameter, with the time-to-peak of diameter (small: 𝑇𝑇𝑃_d_ = 7.96 ± 1.99 s; medium: 8.32 ± 2.11 s; large: 8.80 ± 1.76 s) and flow velocity (small: 𝑇𝑇𝑃_v_ = 7.86 ± 1.67 s; medium: 8.03 ± 1.73 s; large: 8.20 ± 1.57 s), being markedly smaller than that of the corresponding value for the BOLD signal (𝑇𝑇𝑃_*BOLD*_ = 10.58 ± 2.39 s). The details of data processing are given in online Methods.

## 3. Discussion

Over the past decades, BOLD-fMRI has been extensively used as a powerful tool for visualizing and understanding brain activity by detecting changes in blood oxygenation levels in response to neuronal activity. However, BOLD signals are an indirect indicator of brain activity, due to an intricate signal generation process associated with NVC, i.e. neuronal activity triggering a localized microvascular response that is propagated along the vascular tree causing regional adaptations of blood flow and oxygenation levels. Understanding the origin of BOLD signals necessitates investigating the activation-evoked dynamics of neural cells (neurons, astrocytes), vascular cells (pericytes), and the related microscopic hemodynamics. The challenge of multi-channel recording during NVC has been partially addressed by MR-compatible optical recording[19, 20, 22, 25] or MR-compatible electrodes, with the latter allowing for simultaneous recordings of both electroencephalography (EEG) and fMRI[32]. Nevertheless, the spatial resolution of these methods is still limited. In this study, we pioneered the multimodal imaging of Miniscope and fMRI, simultaneously capturing the high-resolution microvascular response and BOLD signals in the S1HL region induced by electrical stimulation of the mouse hind-paw, highlighting the synchronized occurrence of localized vascular dilation, increased blood flow, and BOLD responses.

From the perspective of technical innovation, MR-μScope overcomes the challenging MR compatibility of integrating the sophisticated miniaturized microscope into a high-field MRI. Through novel PCB design and electromagnetic shielding, as well as a customized RF coil, we achieved high SNR during simultaneous dual-modal operation (Fig. 2). For in-vivo studies in mice, the customized animal cradle is equipped with a 4-DOF Miniscope holder and a dual-pin skull holder, allowing accurate positioning of the ROI in focus for individual mice and significantly reducing motion artifacts. The system is also compatible with physiological monitoring and task-based electrical stimulation devices, enhancing its utility for awake animal behavior studies[33]. An attractive feature of the setup is its low cost. Compared with other MRI-optics devices, e.g., the fiber bundle-based system[22, 23] containing expensive optical components, our system combines the advantages of a standalone single-photon excitation Miniscope with compact design and cost-effectiveness.

In the evoked animal experiments, we employed peripheral electrical stimulation at different frequencies and simultaneously recorded microvascular responses and macroscopic BOLD signals using MR-μScope. The results revealed that stimulation at all frequencies led to robust changes in all parameters assessed. Both the microvascular diameter and flow velocity were found to correlate with the amplitude of the BOLD response, consistent with previous multimodal experiments using a similar stimulation paradigm[17, 25]. Our preliminary data indicate that limits to microvascular dilatation, and hence in total hemoglobin are dominating the BOLD signal behaviour, rather than the changes in the blood flow rate as previously reported[31]. Using our MR-μScope will give us a handle to study the underlying substrate of this effect. The amplitude differences of BOLD between 3 Hz and 6 Hz were small and relatively weaker compared to the significantly stronger responses at 9 Hz, for all readouts evaluated (microvascular diameter, flow velocity, BOLD), which is in line with earlier reports on frequency-dependence of stimulus-evoked responses[34, 35].

Our study provides direct evidence that larger vessels exhibit stronger spatial correlation with BOLD fMRI signal amplitude than smaller ones. This validates biophysical predictions that larger blood volume amplifies deoxyhemoglobin’s magnetic susceptibility effect[1, 36]. Prior diffusion-weighted fMRI studies also empirically showed that while oxygen exchange occurs in capillaries, measurable signals originate predominantly from downstream larger vessels[37]. While neuronal/astrocytic vasodilatory signaling is localized to active sites, vascular responses involve vessels > 1 mm away[38, 39]. This occurs via retrograde vasodilation propagation, recently demonstrated in vivo in rat cortex[40]. The endothelial layer guides signal backpropagation to upstream arterioles. FH likely integrates these differentially acting mechanisms[38, 41], aligning with our observation of resting micro-vessel diameter dependence.

As mentioned in the Introduction section, neurological disorder frequently involves vascular disease as well, i.e., changes in the BOLD response may reflect alterations in neuronal processing, in the efficiency of NVC, or a combination of both. Our dual-modal imaging approach enables simultaneous measurements of neural responses (e.g., by probing cellular calcium levels using optical readouts, see below), microvascular metrics alongside the BOLD signal, thus providing potential insights into neurovascular dysfunction within diseased tissue. This is pivotal not only for accurately interpreting dysfunctional neural circuits from fMRI data but also for developing more specific biomarkers that distinguish vascular health from neural integrity.

There are still several limitations in our work. As the first step towards a multi-scale and multi-modal brain imaging platform, the current setup only integrates a single-wavelength detection, resolving pial vessels in the cortex[42]. Standalone multi-wavelength Miniscope is capable of simultaneously detecting changes in Ca2+ and microvascular parameters, providing a more comprehensive perspective for interpreting BOLD signals[43]. Therefore, developing the next-generation multi-wavelength MR-μScope, targeted at multi-object imaging[43], OISI[25], and optogenetics[23], will be highly attractive. Such a cross-scale and multi-modal hybrid system can synergistically reveal the relationships between cellular activities, local blood flow responses, and whole-brain BOLD signals, which will be helpful for an in-depth understanding of the neural vascular coupling mechanism and the source of BOLD signals.

In conclusion, MR-μScope enables simultaneous multi-scale recording of microscopic structure and dynamics with whole-brain structural and functional mapping. Our platform integrates an MR-compatible Miniscope (spatial resolution, 0.7 μm; speed, 60 Hz) with a customized RF coil and an adaptive animal cradle, achieving artifact-free dual-modal imaging with high SNR (fMRI ≥ 15 dB; optics ≥ 26 dB). Simultaneous measurements in mice revealed the spatiotemporal characteristics of microvascular dynamics and global hemodynamic responses. Such information allows establishing a direct link between the microvascular architecture and function and the macroscopic BOLD responses and thus contribute to the understanding of the physiological origin of fMRI contrast, which is of relevance for investigating brain disorders.

## 4. Online Methods

### 4.1 Detailed System Description

#### 1) MR-compatible Miniscope

The MR-compatible Miniscope constitutes a central module of MR-μScope, which is based on the design of the open-source UCLA Miniscope[29]. Major components of the Miniscope include a CMOS imaging sensor PCB, a data acquisition PCB, an illumination module, an optical assembly, and a copper shield casing (Fig. 1b). For MR compatibility, we redesigned the sensor PCB by removing electronic components containing ferromagnetic materials, such as ferrite beads (LQM21PN2R2MC0D, 2.2 μH at 1 Mhz, Murata; BLM18KG601SN1D, 600 Ω, Murata; CIG21L4R7MNE, 4.7 μH at 1 Mhz, Samsung Electro-Mechanics; CBC3225T101MR, 100 μH at 100 kHz, Taiyo Yuden), and replacing metal-sealed oscillators (OSC SG_211SCE 26MHz, Epson) with ceramic ones (OSC 1ZNB26000AB0R 26MHz, Daishinku). In addition, we added an external power supply and replaced the voltage aggregator module (which was a DC-DC converter) with an LDO, thus ensuring MR compatibility without compromising power stability or signal acquisition performance. The optical assembly is comprised of a half-ball lens (#47-269, Edmund, New Jersey, USA), an excitation filter (ET470/40x, Chroma, Vermont, USA), a dichroic mirror (T495lpxr, Chroma, Vermont, USA), a GRIN lens (#64526, Edmund Optics, USA; 1.8 mm diameter), an emission filter (ET525/50m, Chroma, Vermont, USA), and an achromatic lens (#49-923, Edmund, New Jersey, USA). The illumination module adopts a flexible PCB-mounted LED with a peak output power of up to 1.2 mW. The housing of Miniscope is fabricated with polyetheretherketone (PEEK) to ensure magnetic safety, and a customized electromagnetic shield is incorporated around the CMOS sensor to block both the B0 and B1 fields within the MRI scanner. Overall, the Miniscope achieves a spatial resolution of approximately 0.7 μm and an imaging speed of up to 60 Hz.

#### 2) Custom animal cradle and Miniscope holder

We developed an animal cradle to fulfill the spatial constraints inside MRI bore (diameter, 200 mm) and provide essential life holder including anesthesia supply, temperature maintaining/monitoring, etc. (Fig. 1c). The animal cradle is equipped with a gas anesthesia mask which also serves as a holder for the customized coil (Fig. 1e). Additionally, the cradle incorporates a skull holder, a Miniscope holder, a circulating water heating pad, and gas delivery tubes for oxygen/anesthesia mixture. The skull holder secures the mouse brain following cranial window surgery and positions the GRIN lens at the targeted region of interest (ROI). Owing to its bilateral dual-pin design, the skull holder can attach the mouse head to the animal cradle via screw fasteners, with its central ring structure providing anatomical conformity to the skull morphology.

A ring-shaped Miniscope holder was designed to accommodate inter-animal variability in cranial window positioning, facilitating precise four-degree-of-freedom adjustments during installation to optimize FOV (Fig. 1d). The Miniscope holder comprises three components: an upper C-shaped bracket for vertical positioning, a lower C-shaped bracket with slots for lateral adjustments, and a sliding baseplate interfacing with the cradle for anteroposterior positioning. The whole assembly of these components enables wide-angle rotation of the Miniscope. All the holders and their components are secured by PEEK fasteners. The compact design sizes approximately 80 × 80 × 160 mm³, fitting the size of the MRI scanner bore. The animal cradle also contains respiratory gating interfaces and electrical stimulation ports. The incorporated excitation source channels support various behavioral paradigms for task-based experiments, enabling the simultaneous acquisition of whole-brain fMRI signals and microscopic optical imaging in specific brain regions.

#### 3) Custom RF coil and its holder

Our hybrid system is used within a 9.4 T high-field preclinical MRI scanner (BioSpec 94/30, Bruker Biospin GmbH, Germany) with a 114 mm-diameter gradient coil bore and 200 mT/m gradient strength. We developed a customized single-channel transceiver RF surface coil to ensure high-quality MRI data acquisition within limited space (Fig. 1e). The coil sizes 38 × 26 mm² with a 20 × 18 mm² window in the center. The coil tuning and matching circuit contains two ceramic capacitors (12 pF and 15 pF) and two tunable capacitors (both 1-23 pF). Mechanical rods are attached to the tunable capacitors allowing for remote RF frequency adjustment and impedance matching under different loading conditions (Supple. Fig. 2). The MRI FOV covers approximately length × width × depth = 15 × 15 × 10 mm³, warranting coverage of the whole mouse brain. During dual-modal operation, the Miniscope objective lens set was mounted through the coil window and placed on the top of the mouse head. This design significantly improves the SNR of the acquired MRI even with an operating Miniscope positioned closely. The resulting SNRs of structural MRI (T1WI, T2WI) and BOLD-fMRI (EPI) are ≥40 dB and ≥15 dB, respectively, which is comparable with standalone MR imaging performance achieved without Miniscope.

### 4.2 System validation and characterization

#### 1) Phantom preparation

Phantom A was used in the static imaging experiment. An ellipsoidal phantom (lengths on x-/y-/z-axis, 10/10/7 mm) was used for simulating the shape of a mouse brain for system characterization (Fig. 2a). A composite solution of 0.5 g agar powder (originating from Solarbio, China) and 100 g water were utilized to construct the phantom matrix. After 3 min cooling, the phantom surface was covered with a fluorescent tape and a USAF 1951 resolution board (R3L1S4P, Thorlabs, New Jersey, US) for imaging for dual-modal imaging.

Phantom B was used in the dynamic imaging experiment. An ellipsoidal phantom (lengths on x-/y-/z-axis, 10/10/7 mm) with a silicone tube through which fluorescent dye and air passed alternately was used in dynamic MR imaging (Fig. 2h). Compared with Phantom A, the mold of Phantom B has two additional holes at the top, allowing the silicone tube to pass through the phantom along y-axis direction. A composite solution of 0.5 g agar powder and 100 g water was utilized to construct the phantom matrix. After 3 min cooling, the phantom surface was covered with a USAF 1951 resolution board.

#### 2) MRI data acquisition in phantom experiments

The MRI data were acquired using a custom surface coil for both transmission and receiving signals. After a localizer scan for initial positioning, a T1WI-FLASH scan was performed with the following parameters: echo time (TE) = 3 ms, repetition time (TR) = 200 ms, flip angle = 30°, FOV= 30 × 30 mm^2^, matrix size = 256 × 256 (∼117 μm in-plane resolution), slice thickness = 0.5 mm, No. of slices = 21, average = 3. Next, a T2WI-RARE scan was performed with the following parameters: TE = 8.25 ms, TR = 2500 ms, flip angle = 180°, FOV = 30.593 × 20 mm^2^, matrix size = 125 × 125 (∼244 × 160 μm^2^ in-plane resolution), slice thickness = 0.7 mm, No. of slices = 15, average = 2, RARE factor = 8. For both static and dynamic imaging, a gradient-echo EPI scan was performed with the following parameters: TE/TR = 11/1000 ms, flip angle = 63.1°, FOV = 12.8 × 9.2 mm^2^, matrix size = 64 × 46 (∼200 μm in-plane resolution), slice thickness = 0.7 mm, No. of slices = 4, number of repetitions = 180, total scan time = 3 min.

### 4.3 In vivo experiment

#### 1) Animal preparation

The in vivo experiment involves male C57BL/6J mice aged 10-12 weeks. All experimental protocols have been approved by the Institutional Animal Care and Use Committee of ShanghaiTech University. Mice were housed in a vivarium facility under controlled environmental conditions with a 12-hour light/dark cycle, ad libitum access to food and water, a temperature range of 20-26 °C, and relative humidity of 40-60%.

Following induction of anesthesia with 2% isoflurane, mice were secured in a stereotaxic frame under sustained anesthesia (1.5% isoflurane, Supple. Fig. 3a). The scalp was depilated, disinfected, and incised to expose the skull (Supple. Fig. 3b-c). A 2.3 mm diameter cranial window was drilled over the right S1HL region (coordinates: AP -0.9 mm, ML +1.5 mm relative to bregma, Supple. Fig. 3d). A 3D-printed, resin-cast dual-pin skull holder (inner diameter: 7 mm; outer diameter: 8 mm; length: 34 mm; height: 2.68 mm), was affixed to the skull using dental cement. The orientation of the holder was carefully aligned, which was dependent on the cranial window location and the surface curvature of the skull, ensuring full coverage of the surgical site. The holder was filled with 0.5% agarose gel, and then a GRIN lens was implanted 0.36 mm above the target depth relative to the bregma (Supple. Fig. 3e). After 2 minutes for agarose solidification (Supple. Fig. 3f), the holder was sealed with dental cement. Mice were subsequently removed from the stereotaxic apparatus and monitored until full recovery from anesthesia.

#### 2) Miniscope Installation and mouse positioning on the animal cradle

For Miniscope installation, the post-recovery mouse was anesthetized with an intraperitoneal injection of urethane (25%, 1.75 g/kg), administered in three fractional doses at 10-minute intervals[44, 45]. Adequate anesthesia depth was confirmed by the absence or significant reduction of pedal withdrawal reflexes. Fluorescein isothiocyanate-dextran (FITC-Dextran, MW 70,000; 3 mg/mL, 0.1 mL) was administered intravenously via the tail vein. The mouse was secured on the custom animal cradle with the Miniscope and surface coil holders properly installed and adjusted to ensure that the cranial window remained properly positioned underneath the objective to achieve a high-quality optical view (Supple. Fig. 4). An additional intravenous injection of FITC-Dextran (0.3 mL, 3 mg/mL) was administered to further enhance image contrast in microvasculature. Throughout the experiment, core body temperature was maintained at 36-37.5 °C using a feedback-regulated, built-in heating pad (BioSpec 94/30 USR; Bruker Biospin MRI, Germany). The respiratory rate was continuously monitored and maintained within 180-220 breaths per minute.

#### 3) Simultaneous fMRI and optical imaging with electric stimulations

After a localizer scan, a T1WI-FLASH scan was performed to acquire anatomical reference images of the brain using the following setting parameters: TE= 3 ms, TR = 200 ms, flip angle = 30°, FOV = 30 × 30 mm^2^, matrix size = 256 × 256 (∼117 μm in-plane resolution), slice thickness = 0.5 mm, No. of slices = 21, average = 3. Next, a T2WI-RARE scan was taken with the following parameters: TE = 8.25 ms, TR = 2500 ms, flip angle = 180°, FOV = 30.593 × 20 mm^2^, matrix size = 125 × 125 (∼244 × 160 μm^2^ in-plane resolution), slice thickness = 0.7 mm, No. of slices = 15, average = 2, RARE factor = 8. During the stimulation task, sequential gradient-echo EPI scans were performed using the following parameters: TE/TR = 13/2000 ms, flip angle = 63.1°, FOV = 12.8 × 11.2 mm^2^, matrix size = 64 × 56 (∼200 μm in-plane resolution), slice thickness = 0.7 mm, No. of slices = 8, number of repetitions = 205, total scan time = 410 s. The brightness of Miniscope LED was adjusted according to the captured image. The frame rate during acquisition was 30 Hz, and the image size was 752 × 480 pixels. The electrical stimulation was carried out on the left hind-paw using a block pulse design with a current amplitude of 1 mA, a pulse duration of 30/15/10 ms, and frequencies of 3/6/9 Hz, 9%, respectively. A stimulation block lasted 10 s followed by a recovery period of 60 s, which was repeated 5 times. The baseline measurement lasted for 60 s.

### 4.4 Data processing

All data are subjected to standardized preprocessing, quantitative analysis, and cross-modal correlation analysis (Supple. Fig. 5). For fMRI data, the workflow includes preprocessing, and task-based first-level analysis to quantitatively assess significant stimulus-evoked changes in BOLD signals. For Miniscope data, iterative background removal, vessel enhancement and segmentation, and iterative Radon transform algorithms are applied to extract vascular diameter and blood flow velocity. Finally, temporal modeling and correlation analysis are conducted to systematically evaluate the relationship between local microvascular parameters in the right cortex and BOLD-fMRI signals, providing a comprehensive characterization of stimulus-evoked neurovascular coupling.

#### 1) fMRI image processing

During the preprocessing stage, the fMRI images are masked manually to remove the surrounding agar signal that may affect the following data analysis. SPM12 is used for motion correction and slice timing[46]. T1W anatomical images are registered to a mouse brain template with BioImage Suite (bioimagesuite.yale.edu), where mutual information-based nonlinear registration is done iteratively[47]. The resulting nonlinear registration transformation is subsequently applied to the corresponding fMRI data from the same mouse.

After image registration, primary analysis of fMRI data was performed by using a generalized linear model (GLM) in SPM12[46]. The GLM regressors contain a main regressor, which is the convolution of stimuli and canonical hemodynamic response function, and a nuisance regressor accounting for head motion correction. The resulting parameter estimates reflected the activation associated with each experimental condition for each mouse. Voxels with stimulus-evoked activation are deemed significant at a threshold of p < 0.05. For right cortical regions meeting statistical criteria considered as ROI, time series are extracted from the significant voxels. The time series were filtered using a bandpass filter (0.01-0.3 Hz). To obtain the change in BOLD in the right cerebral cortex of the mice, the mean intensity of EPI images within the ROI 60 seconds prior to the stimulation is taken as the baseline to calculate the percentage of the absolute value of the intensity within the ROI relative to the baseline during the entire stimulation period.

#### 2) Miniscope image processing

To enhance fluorescence image quality, we first apply iterative background removal[48]. The convergence criterion of the iteration is defined as a maximum absolute difference between consecutively generated background fields is less than 1%. Afterwards, Frangi filtering is used to further depict the vascular structures[49].

To calculate the vascular diameter, the enhanced images are first binarized to generate a vascular mask. Next, vessel skeletons are extracted from the mask. For each skeleton pixel (𝑥_s_, 𝑦_s_), a distance field d(𝑥_s_, 𝑦_s_) is obtained by measuring the nearest Euclidean distance from (𝑥_s_, 𝑦_s_) to the vascular boundary. The local vascular diameter at pixel (𝑥_s_, 𝑦_s_) was defined as 2d(𝑥_s_, 𝑦_s_) + 1 . A sliding window is applied along the vessel skeleton with overlapping areas assigning the mean value of calculated diameters.

To calculate the blood flow velocity, the vessel skeleton is sampled at an interval of 20 pixels. For each vessel segment, a sliding window with a length of 120 pixels is applied to generate a high SNR kymograph, i.e., time-resolved line-scan profiles based on the orientation of the segment. The movement of non-fluorescent red blood cells appears as dark streaks in a kymograph. Therefore, the blood flow velocity can be derived by measuring the streak slopes in combination with the pixel resolution and frame rate of Miniscope. An open-source iterative Radon transform algorithm[50] is used to determine the velocity for each vessel segment. Similar to diameter analysis, a sliding window is used with overlapping regions assigning mean velocity values. To obtain the relative change of the diameter and blood flow velocity in cerebral microvasculature, the mean diameters and flow velocities 60 seconds prior to the start of the stimulation were taken as the baseline.

#### 3) Data analysis of cross-modality correlation

To investigate the association between local vascular parameters and BOLD-fMRI signals quantitatively, we performed dynamic parameter modeling and cross-modality correlation analysis on the three parameters acquired via MR-μScope.

For each vascular segment, mean diameter and blood flow velocity changes are obtained. A bandpass filter (0.01-3 Hz) is applied to compute the rate of change for diameter and flow velocity of vascular segments. The temporal variation of diameter, blood flow velocity, and BOLD signals within each trial are modeled using a gamma-variant function with nonlinear least squares fitting[51]:

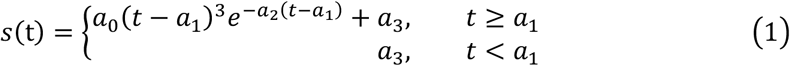

where 𝑎_1_ denotes the onset time and 𝑎_2_ governs the full width at half maximum (FWHM) and 𝑎_3_ the baseline signal. Besides those, the value of AUC of the fitted gamma function is used as a cumulated metric for correlation analysis, which is given by:

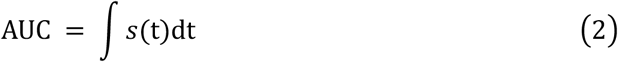

Afterwards, AUC of each trial is filtered based on physiologically plausible onset time, FWHM, and AUC thresholds to ensure signal integrity. Diameter-BOLD and velocity-BOLD relationships are analyzed using linear regression models based on the obtained AUC values. Pearson correlation coefficients are computed to assess statistical significance. For trials with multiple vascular segments, diameter and flow velocity values are averaged to represent the global characteristics of a single trial, which appears as a single dot in Fig. 4-5.

To explore diameter-dependent BOLD correlations, segments are classified into three groups based on mean diameter during stimulation: 15-30 μm, 30-40 μm, and 40-60 μm, which ensures balanced data distribution among groups. The numbers of the effective vascular segments in mouse No. 1-5 under 3/6/9 Hz-stimuli are 48/48/45, 25/31/30, 18/37/34, 29/49/48, and 20/20/18, respectively. Following the filtering procedures described above, 786 valid diameter data points, 398 valid speed data points, and 68 valid BOLD data points were obtained.

## **5.** Acknowledgements

The work is supported by Science and Technology Commission of Shanghai Municipality under Grant 25JS2830300 (Wuwei Ren), National Natural Science Foundation of China under Grants 62105205 (Wuwei Ren), 32170959 (Ning Zhou), 81950410637 (Garth Thompson) and Grant 3210055 (Hui Li).

## Reference

[1] S. Ogawa, T.-M. Lee, A. R. Kay, and D. W. Tank, “Brain magnetic resonance imaging with contrast dependent on blood oxygenation,” proceedings of the National Academy of Sciences, vol. 87, no. 24, pp. 9868–9872, 1990.

[2] A. Lasek-Bal, J. Kidoń, M. Błaszczyszyn, B. Stasiów, and A. Żak, “BOLD fMRI signal in stroke patients and its importance for prognosis in the subacute disease period–Preliminary report,” Neurologia i neurochirurgia polska, vol. 52, no. 3, pp. 341–346, 2018.

[3] N. Franzmeier et al., “Functional brain architecture is associated with the rate of tau accumulation in Alzheimer’s disease,” Nature communications, vol. 11, no. 1, p. 347, 2020.

[4] J. Wang et al., “Perfusion functional MRI reveals cerebral blood flow pattern under psychological stress,” Proceedings of the National Academy of Sciences, vol. 102, no. 49, pp. 17804–17809, 2005.

[5] A. Kawala-Sterniuk et al., “Summary of over fifty years with brain-computer interfaces—a review,” Brain sciences, vol. 11, no. 1, p. 43, 2021.

[6] F. Hyder et al., “Neurovascular and neurometabolic couplings in dynamic calibrated fMRI: transient oxidative neuroenergetics for block-design and event-related paradigms,” Frontiers in neuroenergetics, vol. 2, p. 18, 2010.

[7] T. Takano et al., “Astrocyte-mediated control of cerebral blood flow,” Nature neuroscience, vol. 9, no. 2, pp. 260–267, 2006.

[8] E. C. Peterson, Z. Wang, and G. Britz, “Regulation of cerebral blood flow,” International journal of vascular medicine, vol. 2011, no. 1, p. 823525, 2011.

[9] C. N. Hall et al., “Capillary pericytes regulate cerebral blood flow in health and disease,” Nature, vol. 508, no. 7494, pp. 55–60, 2014.

[10] B. V. Zlokovic, “Neurovascular mechanisms of Alzheimer’s neurodegeneration,” Trends in neurosciences, vol. 28, no. 4, pp. 202–208, 2005.

[11] G. Del Zoppo, “The neurovascular unit in the setting of stroke,” Journal of internal medicine, vol. 267, no. 2, pp. 156–171, 2010.

[12] S. Al-Bachari, R. Vidyasagar, H. C. Emsley, and L. M. Parkes, “Structural and physiological neurovascular changes in idiopathic Parkinson’s disease and its clinical phenotypes,” Journal of Cerebral Blood Flow & Metabolism, vol. 37, no. 10, pp. 3409–3421, 2017.

[13] H. Lütcke et al., “Optical recording of neuronal activity with a genetically-encoded calcium indicator in anesthetized and freely moving mice”, Frontiers in neural circuits, vol. 4, p. 1467, 2010.

[14] W. Mittmann et al., “Two-photon calcium imaging of evoked activity from L5 somatosensory neurons in vivo,” Nature neuroscience, vol. 14, no. 8, pp. 1089–1093, 2011.

[15] Y. B. Sirotin and A. Das, “Anticipatory haemodynamic signals in sensory cortex not predicted by local neuronal activity,” Nature, vol. 457, no. 7228, pp. 475–479, 2009.

[16] A. K. Dunn et al., “Simultaneous imaging of total cerebral hemoglobin concentration, oxygenation, and blood flow during functional activation,” Optics letters, vol. 28, no. 1, pp. 28–30, 2003.

[17] B. G. Sanganahalli, P. Herman, H. Blumenfeld, and F. Hyder, “Oxidative neuroenergetics in event-related paradigms,” Journal of Neuroscience, vol. 29, no. 6, pp. 1707–1718, 2009.

[18] P. Herman, B. G. Sanganahalli, H. Blumenfeld, and F. Hyder, “Cerebral oxygen demand for short-lived and steady-state events,” Journal of neurochemistry, vol. 109, pp. 73–79, 2009.

[19] K. Schulz et al., “Simultaneous BOLD fMRI and fiber-optic calcium recording in rat neocortex,” Nature methods, vol. 9, no. 6, pp. 597–602, 2012.

[20] F. Schlegel et al., “Fiber-optic implant for simultaneous fluorescence-based calcium recordings and BOLD fMRI in mice,” Nature Protocols, vol. 13, p. 840, 03/01 2018, doi: 10.1038/nprot.2018.003.

[21] H.-I. Ioanas, F. Schlegel, Z. Skachokova, A. Schroeter, T. Husak, and M. Rudin, “Hybrid fiber optic-fMRI for multimodal cell-specific recording and manipulation of neural activity in rodents,” Neurophotonics, vol. 9, no. 3, pp. 032206–032206, 2022.

[22] E. M. R. Lake et al., “Simultaneous cortex-wide fluorescence Ca2+ imaging and whole-brain fMRI,” Nature Methods, vol. 17, no. 12, pp. 1262–1271, 2020/12/01 2020, doi: 10.1038/s41592-020-00984-6.

[23] S. Kim et al., “Whole-brain mapping of effective connectivity by fMRI with cortex-wide patterned optogenetics,” Neuron, 2023.

[24] W. Ren et al., “Monitoring mouse brain perfusion with hybrid magnetic resonance optoacoustic tomography,” Biomedical Optics Express, vol. 14, no. 3, pp. 1192–1204, 2023.

[25] R. Bernard, M. Valverde Salzmann, K. Scheffler, and R. Pohmann, “Concurrent intrinsic optical imaging and fMRI at ultra-high field using magnetic field proof optical components,” NMR in Biomedicine, vol. 36, no. 7, p. e4909, 2023.

[26] W. Ren et al., “Dynamic measurement of tumor vascular permeability and perfusion using a hybrid system for simultaneous magnetic resonance and fluorescence imaging,” Molecular imaging and biology, vol. 18, pp. 191–200, 2016.

[27] J. He, I. M. Devonshire, J. E. Mayhew, and N. G. Papadakis, “Simultaneous laser Doppler flowmetry and arterial spin labeling MRI for measurement of functional perfusion changes in the cortex,” Neuroimage, vol. 34, no. 4, pp. 1391–1404, 2007.

[28] M. R. L. Evelyn and J. H. Michael, “Building bridges: simultaneous multimodal neuroimaging approaches for exploring the organization of brain networks,” Neurophotonics, vol. 9, no. 3, p. 032202, 9/1 2022, doi: 10.1117/1.NPh.9.3.032202.

[29] D. J. Cai et al., “A shared neural ensemble links distinct contextual memories encoded close in time,” Nature, vol. 534, no. 7605, pp. 115–118, 2016.

[30] P. Tian et al., “Cortical depth-specific microvascular dilation underlies laminar differences in blood oxygenation level-dependent functional MRI signal,” Proceedings of the National Academy of Sciences, vol. 107, no. 34, pp. 15246–15251, 2010.

[31] R. D. Hoge, J. Atkinson, B. Gill, G. R. Crelier, S. Marrett, and G. B. Pike, “Investigation of BOLD signal dependence on cerebral blood flow and oxygen consumption: the deoxyhemoglobin dilution model,” Magnetic resonance in medicine: An official journal of the international society for magnetic resonance in medicine, vol. 42, no. 5, pp. 849–863, 1999.

[32] M. G. Báez-Yánez, P. Ehses, C. Mirkes, P. S. Tsai, D. Kleinfeld, and K. Scheffler, “The impact of vessel size, orientation and intravascular contribution on the neurovascular fingerprint of BOLD bSSFP fMRI,” NeuroImage, vol. 163, pp. 13–23, 2017.

[33] J. L. Chen, M. L. Andermann, T. Keck, N.-L. Xu, and Y. Ziv, “Imaging neuronal populations in behaving rodents: paradigms for studying neural circuits underlying behavior in the mammalian cortex,” Journal of Neuroscience, vol. 33, no. 45, pp. 17631–17640, 2013.

[34] T. Kim, K. Masamoto, M. Fukuda, A. Vazquez, and S.-G. Kim, “Frequency-dependent neural activity, CBF, and BOLD fMRI to somatosensory stimuli in isoflurane-anesthetized rats,” Neuroimage, vol. 52, no. 1, pp. 224–233, 2010.

[35] S. A. Sheth, M. Nemoto, M. Guiou, M. Walker, N. Pouratian, and A. W. Toga, “Linear and nonlinear relationships between neuronal activity, oxygen metabolism, and hemodynamic responses,” Neuron, vol. 42, no. 2, pp. 347–355, 2004.

[36] J. L. Boxerman et al., “The intravascular contribution to fMRI signal change: Monte Carlo modeling and diffusion-weighted studies in vivo,” Magnetic resonance in medicine, vol. 34, no. 1, pp. 4–10, 1995.

[37] K. Uludağ and P. Blinder, “Linking brain vascular physiology to hemodynamic response in ultra-high field MRI,” Neuroimage, vol. 168, pp. 279–295, 2018.

[38] E. M. Hillman, “Coupling mechanism and significance of the BOLD signal: a status report,” Annu Rev Neurosci, vol. 37, pp. 161–81, 2014, doi: 10.1146/annurev-neuro-071013-014111.

[39] P. Li, Q. Luo, W. Luo, S. Chen, H. Cheng, and S. Zeng, “Spatiotemporal characteristics of cerebral blood volume changes in rat somatosensory cortex evoked by sciatic nerve stimulation and obtained by optical imaging,” J Biomed Opt, vol. 8, no. 4, pp. 629–35, Oct 2003, doi: 10.1117/1.1609199.

[40] B. R. Chen, M. G. Kozberg, M. B. Bouchard, M. A. Shaik, and E. M. Hillman, “A critical role for the vascular endothelium in functional neurovascular coupling in the brain,” J Am Heart Assoc, vol. 3, no. 3, p. e000787, Jun 12 2014, doi: 10.1161/JAHA.114.000787.

[41] D. Attwell, A. M. Buchan, S. Charpak, M. Lauritzen, B. A. Macvicar, and E. A. Newman, “Glial and neuronal control of brain blood flow,” Nature, vol. 468, no. 7321, pp. 232–43, Nov 11 2010, doi: 10.1038/nature09613.

[42] I. Singh, Textbook of Human Neuroanatomy. Jaypee Brothers, 2002.

[43] N. Chen et al., “Simultaneous head-mounted imaging of neural and hemodynamic activities at high spatiotemporal resolution in freely behaving mice,” Science Advances, vol. 11, no. 12, p. eadu1153, 2025.

[44] F. K. Johnson et al., “Amygdala hyper-connectivity in a mouse model of unpredictable early life stress,” Translational psychiatry, vol. 8, no. 1, p. 49, 2018.

[45] K. P. de Arce et al., “Concerted roles of LRRTM1 and SynCAM 1 in organizing prefrontal cortex synapses and cognitive functions,” Nature Communications, vol. 14, no. 1, p. 459, 2023.

[46] W. D. Penny, K. J. Friston, J. T. Ashburner, S. J. Kiebel, and T. E. Nichols, Statistical parametric mapping: the analysis of functional brain images. Elsevier, 2011.

[47] H. Li et al., “Resting state brain networks under inverse agonist versus complete knockout of the cannabinoid receptor 1,” ACS Chemical Neuroscience, vol. 15, no. 8, pp. 1669–1683, 2024.

[48] W. Zhao et al., “Sparse deconvolution improves the resolution of live-cell super-resolution fluorescence microscopy,” Nature biotechnology, vol. 40, no. 4, pp. 606–617, 2022.

[49] A. F. Frangi, W. J. Niessen, K. L. Vincken, and M. A. Viergever, “Multiscale vessel enhancement filtering,” in Medical image computing and computer-assisted intervention—MICCAI’98: first international conference cambridge, MA, USA, october 11–13, 1998 proceedings 1, 1998: Springer, pp. 130–137.

[50] P. Y. Chhatbar and P. Kara, “Improved blood velocity measurements with a hybrid image filtering and iterative Radon transform algorithm,” Frontiers in neuroscience, vol. 7, p. 106, 2013.

[51] J. A. de Zwart et al., “Impulse response timing differences in BOLD and CBV weighted fMRI,” Neuroimage, vol. 181, pp. 292–300, 2018.

[52] Q. Wang et al., “The Allen mouse brain common coordinate framework: a 3D reference atlas,” Cell, vol. 181, no. 4, pp. 936–953. e20, 2020.

